# Genome-Wide Identification of Conditionally Essential Genes Supporting *Streptococcus suis* Growth in Serum and Cerebrospinal Fluid

**DOI:** 10.1101/2025.05.03.652005

**Authors:** Maria Juanpere-Borras, Tiantong Zhao, Jos Boekhorst, Blanca Fernandez-Ciruelos, Rajrita Sanyal, Nissa Arifa, Troy Wagenaar, Peter van Baarlen, Jerry Wells

## Abstract

*Streptococcus suis* is a major cause of sepsis and meningitis in pigs, and zoonosis through the emergence of disease-associated lineages. The ability of *S. suis* to adapt and survive in host environments, such as blood and cerebrospinal fluid (CSF), is important for pathogenesis. Here, we used Tn-Seq coupled with Nanopore sequencing to identify conditionally essential genes for growth of *S. suis* P1/7 in active porcine serum (APS) and CSF derived from choroid plexus organoids (Zhao et al., 2025). Through comparative fitness analyses we identified 33 conditionally essential genes (CEGs) supporting growth in APS and 25 CEGs in CSF, highlighting the importance of pathways involved in amino acid transport, nucleotide metabolism, and cell envelope integrity. Notably, the LiaFSR regulatory system and multiple ABC transporters were important for proliferation. We also identified several genes of unknown function as essential for growth, pointing to previously unrecognized genetic factors involved in *S. suis* adaptation during infection. These findings provide new insights into the genetic requirements for *S. suis* survival in host-like environments and a deeper understanding of its ability to adapt to distinct physiological niches.

## Introduction

*Streptococcus suis* (*S. suis*) is an encapsulated Gram-positive coccus that causes infections of the porcine respiratory tract as well as severe invasive infections such as septicemia, arthritis and meningitis (Feng et al., 2014; Weinert et al., 2015). Diseases caused by *S. suis* results in significant economic losses in the pork industry due to mortality, antimicrobial treatments and the use of autogenous vaccines (Walker, 2024). *S. suis* is also a public health concern for persons encountering diseased pigs or raw pork products due to the high zoonotic potential of specific lineages (Ho et al., 2011; Segura, 2020; Takeuchi et al., 2017). Highest cases of *S. suis* zoonotic disease occur in Southeast Asia, with Vietnam having the highest number of cases (Mai et al., 2008). Currently, 29 virulence-associated serotypes of *S. suis* have been reported, with serotype 2 being the most common cause of invasive disease in swine and human infections (Segura, 2020; Takeuchi et al., 2017).

Asymptomatic carriage of *S. suis* in the upper respiratory, genital, and intestinal tracts of pigs is common (Wertheim et al., 2009). *S. suis* exhibits relatively low invasiveness toward epithelial cells supporting the hypothesis that the main route of entry into the body *is via* the palatine tonsils (Segura et al., 2016; Staats et al., 1997). Immunohistochemistry research shows that *S. suis* can enter deep in the tonsillar crypts, where the surface epithelium becomes a single cell layer thick, facilitating bacterial uptake and translocation (Isabela M. Fernandes de Oliveira, 2023). One hypothesis is that *S. suis* is phagocytosed, but not killed, by specific subsets of tonsillar macrophages, allowing phagocytosed *S. suis* bacteria to replicate and travel through the efferent lymphatics to the bloodstream or directly enter the circulation via the blood vessels in the lymphoid tissue (Segura & Gottschalk, 2002; Williams & Blakemore, 1990). Once in the bloodstream, *S. suis* can cause sepsis, and meningitis upon crossing the blood-brain barrier or the blood-cerebrospinal fluid barrier (Feng et al., 2014; Tram et al., 2021). The precise pathways and molecular mechanisms enabling *S. suis* to infect hosts remain poorly characterized (Segura et al., 2017). This underscores the need for further studies to delineate the genetic and cellular factors underpinning *S. suis* pathogenesis (Segura et al., 2017).

Despite advances in understanding the pathogenesis of *S. suis*, the development of effective vaccines remains challenging. This difficulty arises from the extensive genetic diversity of *S. suis* (Weinert et al., 2015), and the fact that similar virulence functions can be carried out by different genes across pathogenic lineages (Murray et al., 2023; Roodsant et al., 2021). Virulence factors recognised to play important roles in *S. suis* pathogenesis are the cytotoxin suilysin, the capsular polysaccharide, and enolase (Tenenbaum et al., 2016; Zhao et al., 2024). Surface-exposed enolase of *S. suis* hijacks the host plasminogen-plasmin proteolytic system to break down the host extracellular matrix (David Roy, 2014; Tenenbaum et al., 2016; Zhao et al., 2024). Other genes important in pathogenesis are genes involved in stress resistance and metal homeostasis in host niches (Singh Chhatwal Editor, 2013; Tram et al., 2021). To colonise the host, survive and replicate during host infection, *S. suis* must compete with other microorganisms for scarce nutrients and adapt to changes in pH and oxygen levels (Singh Chhatwal Editor, 2013). Bacterial sensory systems play a crucial role in detecting environmental changes, nutrient availability and rapidly regulating gene expression to support replication, which is crucial for pathogenesis and transmission. Hence, a better knowledge and understanding of the genetics regulating metabolic adaptation of *S. suis* to different host niches might lead to better understanding of *S. suis* pathobiology and the development of new therapeutic strategies (Rohmer et al., 2011).

A substantial portion of the *S. suis* genome has hitherto remained uncharacterized, with numerous genes encoding hypothetical proteins with unknown functions, as described in NCBI databases (NC_012925). Consequently, critical genes for infection and pathogenesis may have remained undiscovered. To address this, unbiased genome-wide screening methods are necessary to identify annotated and hypothetical genes necessary for *S. suis* replication and survival in the host.

In this study we used Transposon Sequencing (Tn-seq), a next-generation sequencing technique to identify genes involved in *S. suis* growth and survival in active porcine serum (APS) and cerebrospinal fluid (CSF) extracted from the lumen of iPSC-derived choroid plexus (ChP) organoids (Zhao et al., 2025). CSF within the lumen of 30-40-day-old human ChP organoids closely resembles *in vivo* CSF (Pellegrini et al., 2020). The ChP organoid model offers an alternative approach to extracting CSF from animals or humans. Significant differences in metabolite concentrations have been measured between CSF and serum (Otto et al., 2024). The latter study, conducted on 58 healthy control individuals, showed that methionine, glutamic acid, and glycine were only detected in a small number of CSF samples but were present in nearly all serum samples. Additionally, inosine was exclusively detected in CSF samples, likely due to its critical roles in purine and energy metabolism and in neuroprotection, where it supports neural repair and reduces oxidative stress (Nascimento et al., 2021; Otto et al., 2024). Tn-seq provides a non-biased, systematic approach by utilizing a saturated transposon insertion library (Tn-library) (van Opijnen et al., 2009). The Tn-seq procedure has been recently used in *S. suis* S10, to identify genes involved in pathogenesis. Using an *in vivo* model, bacteria were inoculated intrathecally in pigs, and mutants were recovered from different body fluids (Arenas et al., 2020). In this study, a Tn-library was generated in *S. suis* P1/7, a zoonotic serotype 2 strain (Segura et al., 2017; Singh Chhatwal Editor, 2013), which was cultured under both control and test conditions. By sequencing the flanking regions of transposon insertions and comparing their frequencies between conditions, we identified insertions that either enhanced or impaired bacterial fitness (Van Opijnen et al., 2009). For cost-effective, rapid and accurate in-house sequencing of Tn-libraries, we utilized Oxford Nanopore sequencing technology. To validate the functional role of specific genes, we performed bioassays using chemically defined media (CDM), APS and CSF to assay proliferation capacities of *S. suis* gene-specific deletion mutants.

Our high-throughput screening results revealed the critical role of specific metabolic pathways for *S. suis* P1/7 growth in APS and CSF. Nucleotide metabolism was essential for survival and proliferation in APS while amino acid uptake was critical for proliferation in CSF. We also characterized a so far unknown nucleotide ABC transporter in *S. suis*, essential for purine uptake. Furthermore, our analysis discovered that the LiaFSR three component system and several other uncharacterized genes appear to be involved in *S. suis* pathogenesis. Notably, we demonstrated, for the first time, the feasibility of using CSF extracted from ChP organoids as a model system.

## Materials and Methods

### Bacterial strains and growth conditions

*Streptococcus suis* strain P1/7 was cultured in Todd-Hewitt medium (Thermo Fisher Scientific™, CM0189) supplemented with 0,2% of yeast extract (Thermo Fisher Scientific™, 212750)(THY) at 37 °C with 5% CO_2_. High transformation-efficiency *Escherichia coli* strain Top10 was cultured in LB medium (Merck, 1,102,850,500) at 37 °C with vigorous shaking. Chloramphenicol (Sigma-Aldrich, C0378-25G) was added to the media at a final concentration of 5 μg/ml for *S. suis* and 20 μg/ml for *E. coli*.

### Growth measurements

Growth of *S. suis* wt and deletion mutants in THY were measured by absorbance (OD_600_) every hour for 8 hours. Growth measurements in APS and CSF were performed hourly by making serial dilutions in PBS and plating on agar plates to obtain CFU/ml. For all strains, overnight cultures were first pelleted at 6000 rpm for 5 minutes, and pellets were resuspended in PBS. The appropriate volume of bacterial culture was inoculated into APS or CSF to achieve an OD_600_ of 0.015. For THY growth measurements, bacterial overnight cultures were directly inoculated without prior pelleting and resuspension in PBS.

### Growth curves in chemically defined media

The complete list of basic components and stock solutions used to prepare chemically defined media (CDM) is found in supplementary materials Table S1. For the preparation of 2 ml of CDM, the following constituents were used: 450 µl of CDM buffer, 0.3 mg/ml amino acids, 50 mM of glucose, 0.01 mg/ml of pyruvate, 20 µl of vitamins, 20 µl of metal mixture, 4 µl of manganese, and 2 µl of choline chloride. Nucleobases were added at concentrations ranging from 5 to 100 mg/ml.

Subsequently, 200 µl of CDM was added to each well of a 96-well plate along with 2 µl of THY overnight cultures. OD_600_ readings were obtained using a SpectraMax® M3 Multi-Mode Microplate Reader (Avantor, Radnor, PA, USA) at 37°C.

### Transposon vector cloning

The materials and protocols for constructing the transposon library were provided by Tim Van Opijnen’s lab (Tufts University School of Medicine, Boston, Massachusetts) and modified as described below (Van Opijnen & Camilli, n.d.). *Magellan6* encodes spectinomycin resistance between two inverted repeat sequences (Van Opijnen & Camilli, n.d.). Given that *S. suis* has intrinsic resistance to spectinomycin, we engineered the *Magellan6* plasmid such that the Himar1 *mariner* transposon carried a chloramphenicol resistance gene under the control of the P32 promoter. The plasmid containing the Himar1 *mariner* transposon, *Magellan6*, was purified using the Qiagen Miniprep kit (QIAprep Spin Miniprep Kit, 27106), following the manufacturer’s recommendations. Linearization of the plasmid was performed using the restriction enzymes FastDigest *Sma*I and *Swa*I (Thermo Fisher Scientific™, 10324630 and 15390291). A DNA fragment containing the P32 promoter and chloramphenicol acetyltransferase resistance gene, was amplified by PCR using primers P001 and P002 (Table S2), and the high-fidelity Q5 polymerase (NEB, M0493S), using plasmid pLABTarget as a template (Van Der Els et al., 2018). The DNA fragments were ligated together using Hifi Assembly (NEB, E2621S), to generate plasmid Meg6SS that now encoded Himar1 *mariner* transposon with the desired chloramphenicol resistance gene.

### Generation of Tn-library for *S. suis* P1/7

Genomic DNA (gDNA) and plasmid DNA solutions were concentrated using a SpeedVac (Thermo Fisher Scientific™, RVT5105) until they reached a concentration of at least 350 ng/µl. 36 µl of purified C9T transposase was mixed with 4 µg of gDNA and 4 µg of plasmid and incubated at 30 _o_C for 6 hours. DNA fragments containing the transposon were transformed into *S. suis* using the natural competence method described by Zaccaria et al. (Zaccaria et al., 2014). Briefly, 10 µl aliquots of treated DNA were combined with 5 µl of competence inducer peptide ComS and 100 µl of *S. suis* THY culture at OD_600_ values between 0.035 and 0.05. After 2 hours incubation, aliquots were pooled to a final volume of approximately 900 µl. A 1/10 dilution of 10 µl of the mixture was plated on THY chloramphenicol agar plates for CFU counting and the remaining 900 μl centrifuged at 6000 rpm for 5 min. Then, 800 µl of supernatant was removed and the pellet resuspended in the remaining 100 µl before plating on THY agar plates containing 5 µg/ml chloramphenicol. This process was repeated four times, resulting in 4 libraries with colony counts of approximately 10K, 1.5K, 3.3K, and <1K, respectively.

### Generation of choroid plexus organoids and extraction of cerebrospinal fluid

Six-well plates (Corning, 07-200-83) were pre-coated with Vitronectin (STEMCELL, 07180) in CellAdhere™ Dilution Buffer (STEMCELL, 07183) at room temperature for 1 hour. Human iPSC line EDi002-A (EBiSC™) were maintained on vitronectin-coated 6-well plates in mTeSR1 (STEMCELL, 85857). Media were changed daily, and cells were passaged once a week. ChP organoids were generated using the STEMdiff™ Choroid Plexus Organoid Differentiation Kit (STEMCELL, 100-0824) and the STEMdiff™ Choroid Plexus Organoid Maturation Kit (STEMCELL, 100-0825), following previous protocols (Pellegrini et al., 2020b) with minor modifications. Briefly, iPSC were dissociated into single-cell suspensions using Accutase (STEMCELL, 07920). On day 1, 1 × 10^5^ cells were seeded into a well of Corning^®^ 96-well round-bottom ultra-low attachment microplate (Corning, 7007) in 100 µL of embryoid body (EB) Formation Medium and 10 µM Y-27632 (ROCK inhibitor; STEMCELL, 72302). On day 2 and day 4, fresh 100 µL of EB Formation Medium was added to each well. EBs with diameter ranging between 400 and 600 µm were typically observed on day 5 at which time EB Formation Medium was replaced with 200 µL/well of Induction Medium. On day 7, each EB was embedded in 15 µL of Matrigel^®^ (Corning, 734-1101) dropwise on sheets of parafilm and incubated at 37°C for 30 min to polymerize Matrigel^®^ (16 EB per sheet of parafilm). The sheets of parafilm were each positioned above one well of a 6-well ultra-low adherent plate (STEMCELL, 100-0083) using sterile forceps. All 16 Matrigel^®^ droplets were gently washed off sheets and put into one well using 3 mL of Expansion Medium. Each 6-well plate was shaken back and forth three times to ensure even distribution of EB and incubated at 37°C for 3 days. On day 10 the Expansion Medium was carefully replaced with 3 mL/well of Choroid Plexus Differentiation Medium, and plates were placed on a platform rotator (Fisherbrand™, 15504080) in the incubator. On day 13, Choroid Plexus Differentiation Medium was refreshed. From day 15, Choroid Plexus Differentiation Medium was replaced with 3 mL/well Maturation Medium and renewed every 3 days. By day 30, ChP organoids epithelia resemble cyst-like structures filled with CSF-like fluid. ChP organoids between day 30 to day 40 were used to harvest CSF from the organoid lumen using a 0.30 × 12 mm BL/LB needle attached to a 1 mL syringe.

### Transposon library screening

For APS screening, 60 µl of *S. suis* Tn-library were inoculated into 10 ml pre-warmed THY broth or APS (Thermo Fisher Scientific™, 26250084), with two biological replicates for each condition and incubated for 5 hours at 37 °C with 5% CO_2_. For CSF screening, 250 µl of Tn-library stock was thawed in 10 ml of THY medium and cultured for 3.5 hours until it reached concentration of 5 × 10^8^ CFU/ml, and then centrifuged and resuspended in 10 ml of PBS. Then 100 µL of the enriched Tn-library in PBS (Fisher Scientific, 10769033) was inoculated in 10 ml of THY and 10 ml of CSF in triplicates and cultured at 37 °C with 5% CO_2_ until cultures reached an OD_600_ of 0.6. Cultures were centrifuged at 4000 rpm for 10 min, the supernatant discarded, and the bacterial pellets stored at -20 degrees until further use. DNA was extracted from bacterial pellets using DNeasy PowerSoil kit (Qiagen, 47016) following the manufacturer’s recommendations.

### Sample preparation

After library growth under the different conditions (APS, CSF and THY), bacterial DNA was isolated and processed as described by Van Opijnen & Camilli (Van Opijnen & Camilli, 2010). Briefly, approximately 2 µg of genomic DNA was digested with the *MmeI* restriction enzyme (NEB, R0637L) and dephosphorylated using Quick CIP (calf intestinal alkaline phosphatase) (NEB, M0525S) to avoid re-ligation. The oligonucleotide primers incorporating the adapter sequences (P003 and P004, Table S2), were annealed by mixing equimolar concentrations of each primer in water in 1.5 ml Eppendorf tubes, heating the tubes to 96°C for 3 minutes, and gradually cooling the tubes to room temperature. Phosphorylated adapters were ligated to the overhangs generated by digestion of bacterial DNA with *MmeI*, using T4 DNA ligase (NEB, M0202S). After adapters ligation, transposon-containing fragments of bacterial DNA were amplified by PCR. The forward primer annealed to the inverted repeat sequence that is present at each end of the transposon, while the reverse primer annealed to ligated adapter (P005 and P006, Table S2). The PCR reaction included 2 µl of the ligation mix and High-fidelity Q5 polymerase (NEB, M0493S) for 35 cycles. Total volumes of PCR reactions were loaded onto 0.8% agarose gel, and bands corresponding to 300 bp that were expected to contain transposon-enriched fragments were excised and purified using Invisorb fragment cleanup kit (Invitek molecular, 1020300300) (Figure 1).

**Figure 1.**
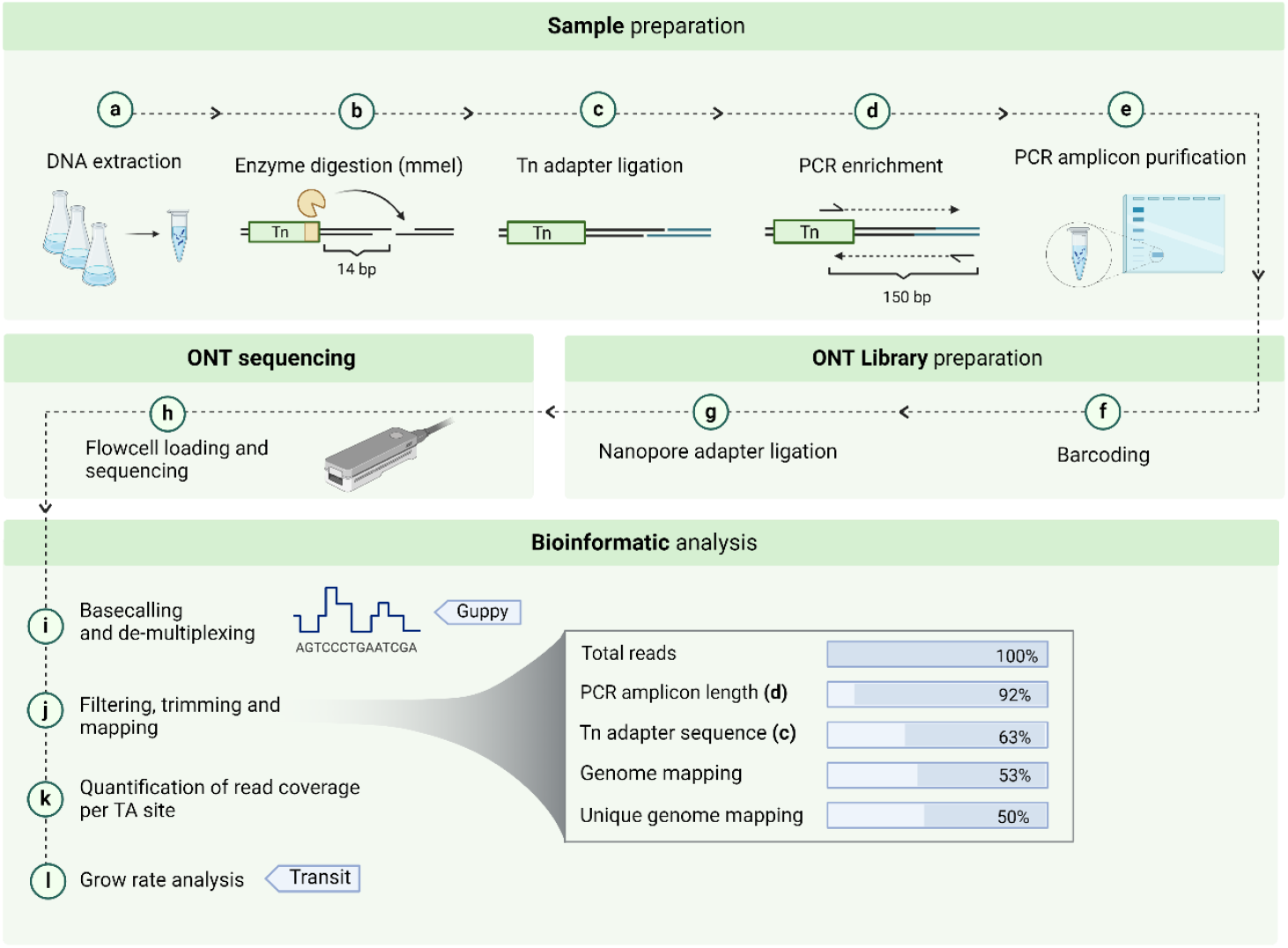
Workflow for Tn-Seq sample processing, sequencing, and bioinformatic analysis. (a-e) Sample preparation begins with DNA extraction, followed by enzyme digestion with MmeI, Tn adapter ligation, PCR enrichment, and amplicon purification. (f-h) ONT Library sequencing. (i-l) in-silico processing of sequencing output and data analysis. (Steps j and k were specifically developed for this study)

### Library sequencing with Nanopore

PCR amplicons were prepared and barcoded for sequencing following the manufacturer’s instructions using the native barcoding kit (ONT, SQK-NBD112.24), the ligation sequencing kit (ONT, SQK-LSK112) kit and flow cells (ONT, FLO-MIN106D) on MinION Mk1c and Mk1b (ONT, Oxford, United Kingdom) sequencing devices; device outputs were handled using MinKNOW software (version 23.07.15). Sequencing ran until all reads were sequenced (approximately 24 hours). The resulting FAST5 files, containing raw nanopore data of each sample, were converted to FASTQ files via basecalling (Guppy version. 7.1.4).

### Analysis of Tn-seq data

For nanopore reads sequence analysis (Fig. 1), we filtered reads to include those within the expected size range of our PCR amplicon, roughly 150 to 180 base pairs (bp). Subsequently, we conducted a two-step screening process for the 20 bp of transposon sequence upstream of the transposon insertion site so that only reads containing the forward transposon (Tn) sequence (ACTTATCATCCAACCTGTTA) or the reverse Tn sequence (TAACAGGTTGGATGATAAGT) were retained for further analysis. For the filtered reads, only the 14 bp immediately adjacent to the respective forward or reverse Tn sequences were selected for alignment to the *S. suis* P1/7 reference genome using Bowtie2. Only reads with 100% identity to a single gene fragment were retained; reads that mapped to more than one gene in the genome were discarded. The number of reads per TA site was quantified, and the data was compiled into a Wiggle (WIG) file. TA sites with zero insertions (no matches) were also included in the WIG file during harmonization to ensure the data includes all TA sites per gene, for calculation and comparison during the fold-change analysis. The harmonized wig files from each replicate of both the control and test conditions were used as input into Transit (version 3.2.7) for growth rate analysis using the Transit “resampling test” module with default parameters. Genes that exhibited a log fold change >1 and achieved statistical significance (adj. p-value < 0.05) were compiled into a final list (Table 1). The set of filtered genes and encoded proteins were annotated by retrieving the corresponding gene descriptions and biological process annotations from NCBI and KEGG databases, respectively.

**Table 1.** List of conditionally essential genes, identified by Tn-seq, that were conditionally essential to support S. suis growth in APS and CSF compared to THY (adj. p-vale < 0.05; Log_2_FC < -1). Genes are ordered from lowest to highest FC; genes belonging to the same operon are marked with the same symbol in the ‘Operons’ column.

| Genes identified by tn-seq in APS |  |  |  |  |  |
| --- | --- | --- | --- | --- | --- |
| Locus tag | Name | Product | Main biological process | Operons | Log<sub>2</sub>FC |
| SSU_RS08850 | - | adenylosuccinate synthase | Purine metabolism |  | -3,19 |
| SSU_RS00280 | *purB* | adenylosuccinate lyase | Purine metabolism |  | -2,84 |
| SSU_RS06600 | - | ABC transporter permease | Transporter | □ | -2,28 |
| SSU_RS02595 | *ppc* | phosphoenolpyruvate carboxylase | Carbon metabolism |  | -1,86 |
| SSU_RS05280 | *ptsP* | phosphoenolpyruvate--protein phosphotransferase | Transporter |  | -1,85 |
| SSU_RS06605 | *trpX* | tryptophan ABC transporter substrate-binding protein | Transporter | □ | -1,84 |
| SSU_RS04185 | *guaA* | glutamine-hydrolyzing GMP synthase | Purine metabolism |  | -1,77 |
| SSU_RS06595 | - | ABC transporter ATP-binding protein | Transporter | □ | -1,73 |
| SSU_RS06820 | - | bifunctional oligoribonuclease/PAP phosphatase NrnA | Sulfur metabolism |  | -1,65 |
| SSU_RS09155 | - | hypothetical protein | NI |  | -1,65 |
| SSU_RS09655 | - | LacI family DNA-binding transcriptional regulator | Transcription factor |  | -1,58 |
| SSU_RS06385 | - | acyl-ACP thioesterase domain-containing protein | NI |  | -1,56 |
| SSU_RS06740 | - | hypothetical protein | NI |  | -1,53 |
| SSU_RS01575 | - | Fur family transcriptional regulator | Transcriptional factor |  | -1,43 |
| SSU_RS04755 | - | BMP family protein | Transporter | ● | -1,34 |
| SSU_RS07155 | - | YtxH domain-containing protein | NI |  | -1,34 |
| SSU_RS05100 | - | mannonate dehydratase | Carbohydrate metabolism |  | -1,29 |
| SSU_RS02715 | - | amino acid ABC transporter substrate-binding protein | Transporter | ◇ | -1,26 |
| SSU_RS02080 | *liaF* | cell wall-active antibiotics response protein LiaF | Signal transduction |  | -1,21 |
| SSU_RS04655 | *pta* | phosphate acetyltransferase | Carbon metabolism |  | -1,18 |
| SSU_RS07195 | - | hypothetical protein | NI |  | -1,18 |
| SSU_RS06140 | *tpx* | thiol peroxidase | Oxidoreductases |  | -1,17 |
| SSU_RS09860 | *guaB* | IMP dehydrogenase | Purine metabolism |  | -1,17 |
| SSU_RS04750 | - | ABC transporter ATP-binding protein | Transporter | ● | -1,15 |
| SSU_RS04745 | - | ABC transporter permease | Transporter | ● | -1,14 |
| SSU_RS02750 | - | DegV family protein | NI |  | -1,11 |
| SSU_RS05325 | - | LacI family DNA-binding transcriptional regulator | Transcriptional factor |  | -1,11 |
| SSU_RS06410 | - | arsenate reductase | Oxidoreductases |  | -1,11 |
| SSU_RS06645 | - | lactonase family protein | Carbon metabolism |  | -1,1 |
| SSU_RS05710 | - | glucosaminidase domain-containing protein |  |  | -1,05 |
| SSU_RS04420 | *pyrE* | orotate phosphoribosyltransferase | Pyrimidine metabolism |  | -1,02 |
| SSU_RS04510 | - | ABC transporter substrate-binding protein/permease | Transporter | ◇ | -1,02 |

| SSU_RS06155 | *tehB* | SAM-dependent methyltransferase TehB | Transferases |  | **-1,01** |
| --- | --- | --- | --- | --- | --- |
| **Genes identified by tn-seq in CSF** |  |  |  |  |  |
| Locus tag | Name | Product | Main biological process | Operons | Log<sub>2</sub>FC |
| SSU_RS02635 | - | hypothetical protein | NI |  | **-3.22** |
| SSU_RS06605 | *trpX* | tryptophan ABC transporter substrate-binding protein | Transporter | □ | **-2.97** |
| SSU_RS06600 | - | ABC transporter permease | Transporter | □ | **-2.88** |
| SSU_RS01560 | - | hypothetical protein | NI |  | **-2.78** |
| SSU_RS06595 | - | ABC transporter ATP-binding protein | Transporter | □ | **-2.63** |
| SSU_RS00910 | *pgk* | Phosphoglycerate kinase | Carbohydrate metabolism |  | **-2.61** |
| SSU_RS02500 | - | hypothetical protein | NI |  | **-2.61** |
| SSU_RS07925 | *glyQ* | Glycine--tRNA ligase alpha subunit | Translation |  | **-2.55** |
| SSU_RS04515 | *artM* | Arginine transport ATP-binding protein ArtM | Transporter |  | **-2.54** |
| SSU_RS06355 | *mscL* | Large-conductance mechanosensitive channel | Signaling and cellular processes |  | **-2.41** |
| SSU_RS03420 | *srlR* | Glucitol operon repressor | Transcriptional factor |  | **-2.17** |
| SSU_RS04835 | *pstB3_1* | Phosphate import ATP-binding protein PstB | Transporter |  | **-2.13** |
| SSU_RS08105 | *fabH* | 3-oxoacyl-[acyl-carrier-protein] synthase 3 | Lipid metabolism |  | **-1.85** |
| SSU_RS07155 | - | hypothetical protein | NI |  | **-1.43** |
| SSU_RS06420 | *rpiA* | Ribose-5-phosphate isomerase A | Carbohydrate metabolism |  | **-1.42** |
| SSU_RS04600 | *lacR_2* | Lactose phosphotransferase system repressor | Transcriptional factor |  | **-1.38** |
| SSU_RS02705 | *yecS_2* | L-cystine transport system permease protein YecS | Transporter | ◇ | **-1.3** |
| SSU_RS08470 | *ilvH* | Putative acetolactate synthase small subunit | Carbohydrate metabolism |  | **-1.3** |
| SSU_RS02715 | *fliY* | L-cystine-binding protein FliY | Transporter | ◇ | **-1.14** |
| SSU_RS03250 | - | Phosphoglycerate dehydrogenase | Energy/Amino acid metabolism |  | **-1.1** |
| SSU_RS02710 | *glnQ_2* | Glutamine transport ATP-binding protein GlnQ | Transporter | ◇ | **-1.08** |
| SSU_RS07955 | *metQ* | putative D-methionine-binding lipoprotein MetQ | Transporter |  | **-1.03** |
| SSU_RS07855 | - | hypothetical protein | NI |  | **-1.02** |
| SSU_RS03205 | - | hypothetical protein | NI |  | **-1.01** |
| SSU_RS05450 | - | hypothetical protein | NI |  | **-1** |

### Gene deletion mutants

In frame deletion mutants were constructed using CRISPR/Cas9 based technology essentially as described by Gussak et al (Gussak et al., 2023). Briefly, mutants were generated by transforming *S. suis* P1/7 with a gRNA-Cas9 co-expression plasmid (pSStarget) and a linear DNA repair template. Upon transformation, Cas9 endonuclease was expressed and directed by the co-expressed guide RNA to specific target gene, where Cas9 induced a double-stranded cut that was repaired by the *S. suis* homologous recombination machinery using the introduced DNA repair template. Resulting colonies were screened for gene-specific deletion mutants and site-specific mutant strains were cured from expression plasmids.

Three different 20 bp guide RNAs were designed for each target gene. Single-stranded oligonucleotides were designed using Benchling software, and synthesized by IDT Technologies. Each primer included a 4 bp overhang compatible with the overhangs of the linearized pSStarget plasmid (P007 to P048, Table S2). Equimolar concentrations of complementary primers for each guide were mixed with annealing buffer (10 mM Tris, pH 7.5, 50 mM NaCl, 1 mM EDTA) and annealed in a thermocycler (Bio-Rad, Hercules, CA, USA) (5 min at 95°C and gradually cooling to 25°C at 1 °C/min). The empty pSStarget plasmid was linearized using the BsaI enzyme (NEB, R3733S) and purified Invisorb Fragment CleanUp kit (Invitek, 1020300300). The annealed guide and digested plasmid were ligated using T4 DNA ligase (NEB, M0202S) for 1 hour at room temperature and subsequently incubated overnight at 4 °C. The ligation mixture was transformed into chemically competent *E. coli* Top10 cells and plated on LB agar plates containing 10 μg/ml of chloramphenicol, followed by overnight incubation at 37 °C. Colonies were screened for correct plasmid constructs using primers P077 and P078 (Table S2), and plasmids and guide were extracted from these colonies using the QIAprep Spin Miniprep Kit (Qiagen).

The repair template was constructed by amplifying approximately 1000 bp upstream and downstream of the target gene using primers (P049 to P076, Table S2). These primers were designed to include approximately the first and last 30 bp of the target gene, ensuring that the entire gene was not deleted to avoid potential polar effects on downstream genes. PCR products were subsequently ligated through Splicing by Overlap Extension (SOE) PCR using external primers, HA1_fwd_X and HA2_rev_X (Table S1). After each PCR step, the PCR products were purified using the Invisorb Fragment CleanUp kit (Invitek, 1020300300).

gRNA-Cas9 co-expression plasmids and repair templates were transformed into *S. suis* using the natural competence method described by Zaccaria et al. (Zaccaria et al., 2014). Briefly, 250 µl of an overnight culture of *S. suis* P1/7 wt were inoculated into 10 ml of THY broth and grown until the OD_600_ reached between 0.035 and 0.058. A 100 µl aliquot of *S. suis* P1/7 culture was combined with 5 µl of competence-inducing peptide ComS, 200–500 ng of plasmid DNA, and 1 µg of repair template. The mixture was incubated at 37 °C with 5% CO_2_ for 2 hours and plated on THY agar plates containing 5 µg/ml of chloramphenicol. Colonies with the correct gene deletion were identified using PCR and external primers HA1_fwd_X and HA2_rev_X (Table S2), and these colonies were cured from plasmids by performing two consecutive overnight passages in THY medium without chloramphenicol.

### In silico amino acid sequence alignments and protein structure predictions

Amino acid sequences were obtained from the *S. suis* P1/7 genome annotation in NCBI (nucleotide ID NC_012925) for SSU_RS04755 and from Uniprot for PnrA (A0A0H2UPF3) and TmpC (P29724). Multiple sequence alignment of the amino acid sequences of SSU_RS04755, PnrA and TmpC were performed using the Clustal Omega algorithm integrated within Jalview (version 2.11.3.3) software, using default parameters. Structure prediction of SSU_RS04755 was performed using the SWISS-MODEL server (https://swissmodel.expasy.org). Structure alignment of the predicted protein encoded by SSU_RS04755 from *S. suis* and PnrA was performed using PyMOL (version 3.0.3).

### Identification of Promoter Regions with Putative LiaR Binding Boxes

Identification of promoter regions potentially containing a LiaR binding box was performed using the FIMO tool from the MEME suite (https://meme-suite.org/meme/tools/fimo) (Grant et al., 2011), utilizing the LiaR binding box motif identified from *Bacillus subtilis* as input (Jordan et al., 2006). The resulting list of genes was manually curated to retain only sequences with p-value < 0.05 and located within 200 bp upstream of a gene. This analysis identified 48 genes with a putative LiaR binding box in their promoter regions (Table S3).

### Analysis of Differential Gene Expression by qPCR

*S. suis* P1/7 and *S. suis* Δ*liaR* mutant were grown to exponential phase (approximately OD_600_ 0.3) in THY broth at 37 °C with 0.5% CO_2_. A 10 ml aliquot was pelleted by centrifugation at 4000 rpm for 10 minutes, the supernatant was discarded, and the pellet snap-frozen in liquid nitrogen before being stored at -80 °C overnight. RNA was extracted from the pelleted cells using the RNeasy Mini Kit (Qiagen, 74104) following the manufacturer’s instructions with specific modifications. Pellets were resuspended in 700 µl of RLT buffer containing 0.1% β-mercafigptoethanol and transferred to lysing matrix B 2 ml tubes (MP Biomedicals, 6911100). Bacterial cells were lysed using a FastPrep-24™ 5G bead beating grinder and lysis system (MP Biomedicals, Solon, OH, USA) with settings; 4.0 m/sec, All-MetalQuickprep adapter, 40 seconds. The tubes were centrifuged for 1 minute at 10,000 rpm, and the supernatant was transferred to a clean Eppendorf tube. Subsequent steps were carried out using the manufacturer’s protocol. Final RNA concentrations were measured using the Qubit RNA Broad Range Kit (Thermo Fisher Scientific™, Q10211) and a Qubit 4 fluorometer (Thermo Fisher Scientific™, Waltham, MA, USA). The Quantitect Reverse Transcriptase Kit (Qiagen, 205311) was used for DNA deletion and cDNA synthesis, with 500 ng of RNA as input. For differential gene expression analysis, 96-well white PCR plates (Bio-Rad, ML9651) and GoTaq qPCR Master Mix (Promega, A6002) were used in a CFX96 Real-Time PCR System (Bio-Rad, Hercules, CA, USA). Primers for each gene are listed in Table S1 (from P079 to P086). Gene expression was calculated using the 2-ΔΔCt method relative to reference gene *gyrA*. Independent experiments were performed in triplicate with three biological replicates each.

## Results

### Obtaining *in vitro* Tn-libraries for *S. suis* P1/7

The transposon library was constructed using a modification of the Tn-Seq protocol optimized for *S. pneumoniae* described by Tim van Opijnen and colleagues (Van Opijnen et al., 2009), to optimize the protocol for *S. suis* P1/7 (see Methods). Using this optimized protocol, we obtained four libraries that were combined to obtain 1 single library of approximately 15K mutants.

### Development of a nanopore sequencing protocol for sequencing of the Tn-libraries

After culturing the Tn-library in test (APS or CSF) and control (laboratory culture medium THY) media, DNA was extracted and processed as outlined in Methods section to obtain a PCR amplicon suitable for sequencing. We opted for nanopore PCR amplicons sequencing to have an in-house method for amplicon sequencing and analysis that could be optimized when appropriate. As it was the first time nanopore technology had been used for this purpose, we designed an in-house pipeline for processing and analyzing the sequenced nanopore reads (Figure 1), which is available at (https://github.com/MariaJuanpereBorras/Nanopore-TnSeq-Pipeline). For each sample, i.e. each Tn-library aliquot cultured in test or control media, nanopore sequencing generated between 0.7 and 5.4 million reads, which were filtered to remove fragments larger or smaller than the expected length of the amplicon (around 180 bp), resulting in an average discard rate of 5%. In the subsequent step, reads were selected based on presence of the transposon sequence (underlined region of P005, Table S2); 64% of the total reads were retained during this phase. After this step, reads were trimmed to retain only the 14 base pairs immediately adjacent to the transposon end sequence. When these 14 bp inserts were mapped to the *S. suis* P1/7 genome (NCBI accession no. NC_012925), 55% of the *de novo* sequenced reads had a perfect match and 51% had a unique perfect match; the remaining 4% were discarded because they did not map to a unique genomic location. After running the harmonized wig files in Transit (version 3.2.7) (DeJesus et al., 2015), a final list detailing the increased or decreased log2 fold change (logFC) for each gene in the test condition compared to the control was generated. Filtering the data on negative logFC (< -1) and adjusted p-value (< 0,05), we obtained a final list of 33 conditionally essential genes (CEGs) for *S. suis* P1/7 grown in APS and 25 CEGs in CSF, both compared to growth in THY control medium (Table 1).

### Genes identified by Tn-seq exhibiting fitness impact in *S. suis* growth in APS compared to THY

The transposon library was cultivated for approximately four hours in THY or APS and Tn-seq was used to identify conditionally essential genes (CEG) that support growth of *S. suis* in APS. Out of the 33 CEG (Table 1), 7 (21%) did not have annotations in NCBI nor KEGG database. Among the annotated genes, 10 (30%) were predicted to be involved in metabolism, 5 (15%) in nucleotide metabolism and 8 genes (24%) in substrate transport. Thus ‘metabolism’ and ‘substrate transport’ were the most abundant annotations of CEGs linked to growth of *S. suis* in APS (Figure 2). Furthermore, 3 genes were annotated as transcription factors, 2 as oxidoreductases, 1 as involved in signal transduction, and 1 as methyltransferase.

**Figure 2.**
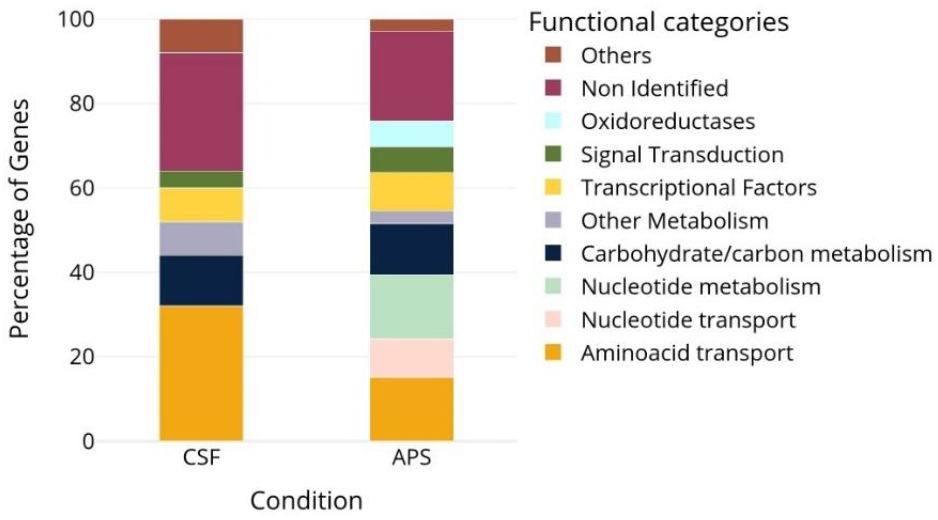
Stacked bar chart showing the percentage of CEGs (logFC < –1, adjusted p-value < 0.05) classified into functional categories for CSF and APS samples.

Of the 10 genes annotated as substrate transporters, 3 genes were predicted to belong to a tryptophan ABC transporter, 3 genes to an amino acid ABC transporter, and 1 gene (gene symbol *ptsP*) as a component of a sugar transporter. The remaining 3 genes were part of a single operon with unknown function. In this operon, a predicted substrate-binding lipoprotein (SSU_RS04755) is highly conserved across *S. suis* strains, including strains from serotype 2 and 9 (Gómez-Gascón et al., 2018). This protein is also part of the *S. suis* secretome and has been studied as a potential vaccine candidate targeting *S. suis (Gómez-Gascón et al., 2018; Weiße et al., 2021).* An amino acid sequence homology search using NCBI blastp tool revealed a high identity (> 60%) with ABC transporters described as nucleoside transporters from *Streptococcus pneumoniae* (SPD_0739) and *Streptococcus agalactiae* (GBS0942) (Franza et al., 2021; Jahn et al., 2022).

### Genes identified by Tn-seq contributing to *S. suis* growth in CSF compared to THY

To explore whether nucleotide metabolism and transport pathways also play a critical role in *S. suis* growth in cerebrospinal fluid (CSF) we performed a Tn-seq screen comparing growth in CSF and THY. In contrast to APS, no genes involved in nucleotide metabolism or transport were conditionally essential for *S. suis* growth in CSF. *S. suis* CEGs supporting growth in CSF included 8 genes (32%) annotated with roles in amino acid transport, 7 genes (28%) annotated as hypothetical proteins, 3 genes annotated with roles in carbohydrate metabolism, and 2 genes annotated as transcriptional regulators (Table 1, Figure 2).

We found 8 CEGs that supported growth in both CSF and APS. The three genes that are predicted to form a tryptophan ABC transporter were identified as conditionally essential in both media (Table 2). Additionally, SSU_RS07195, annotated as hypothetical protein, and SSU_RS02715, annotated as cystine-binding lipoprotein were also identified as conditionally essential for growth in CSF and APS. Transposon insertions in *liaF*, a membrane protein, and the hypothetical protein SSU_RS07195, were associated with increased proliferation during growth in CSF compared to THY, whereas insertions in the same genes (*liaF* and SSU_RS07195), were associated with reduced proliferation in APS compared to THY. Lastly, transposon insertions in the phosphate importer *pstB3_1* were associated with increased proliferation in APS but reduced proliferation in CSF.

**Table 2.**
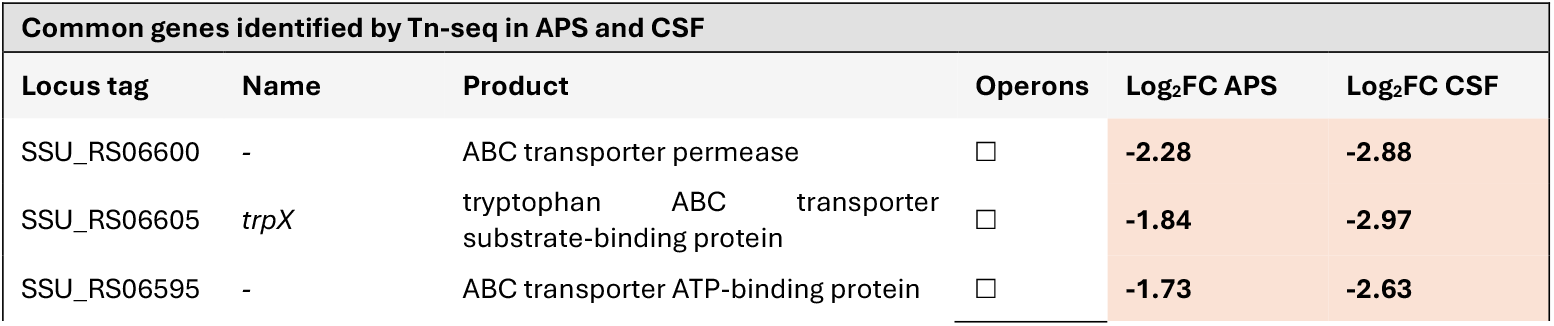
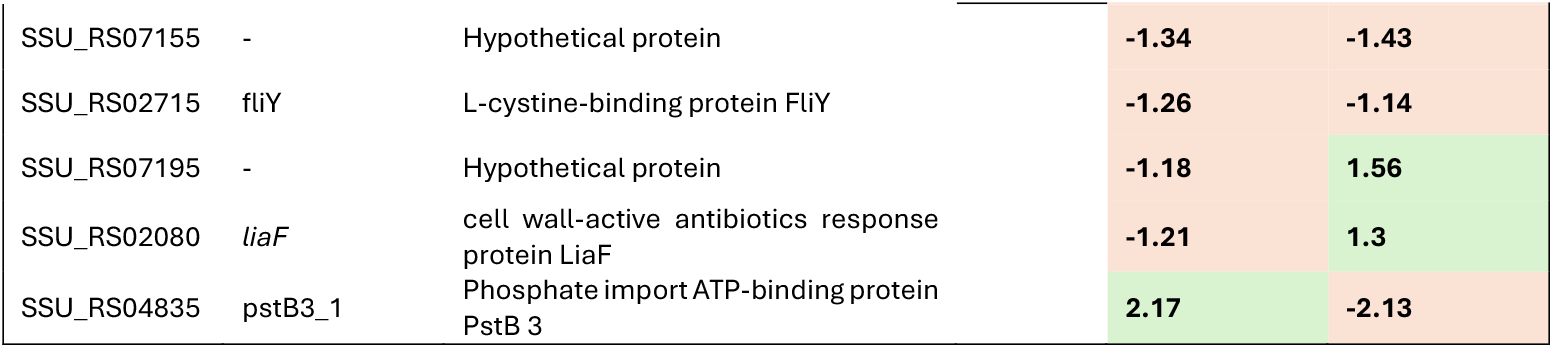
Shared genes identified by Tn-seq analysis of S. suis P1/7 in APS and CSF media (adj. p-vale < 0.05; Log_2_FC < - 1; Log_2_FC > 1). genes belonging to the same operon are marked with the same symbol in the ‘Operons’ column. Genes with decreased fitness (negative Log_2_FC) are highlighted in orange, and genes with increased fitness (positive Log_2_FC) are highlighted in green.

### Validating Tn-Seq results using In-Frame Deletion Mutants

To validate the Tn-seq results, we selected 5 CEGs identified in APS (*purA*, SSU_RS04755, *liaF*, SSU_RS09155 and SSU_RS07155; Table 1) to make in-frame deletions in *S. suis* strain P1/7. Given that genes related to nucleotide metabolism and transport were the most represented among the CEGs (Figure 2), we hypothesized that nucleotide availability was a limiting factor for *S. suis* growth in serum. We chose to generate a *purA* (SSU_RS08850) deletion mutant strain because (i) *purA* was annotated as playing a role in nucleotide metabolism; (ii) showed the highest fold change difference (−3.19) in the list of CEGs, and (iii) was reported as a CEG for *S. suis* survival in pigs in previous Tn-seq studies (Arenas et al., 2020). SSU_RS04755 was selected because it is within a predicted nucleoside ABC transporter operon. Additionally, we selected the predicted *liaF* (SSU_RS02080) gene, annotated as participating in signal transduction. We had found that *liaF* was conditionally essential when *S. suis* was cultured in APS, with a negative logFC, and its inactivation increased fitness in CSF resulting in positive logFC (see above). This suggests a differential role for *liaF* in growth under these two culture conditions. In various Gram-positive bacteria, *liaF* is part of a three-component system, along with a histidine kinase (*liaS*) and a transcription regulator (*liaR*)(Wolf et al., 2010). *LiaFSR* is involved in virulence and antibiotic resistance in streptococci (Suntharalingam et al., 2009; Vega et al., 2024), but it has not been studied in *S. suis*. We were unable to obtain a viable *liaF* deletion mutant. Downstream of *liaF* are *liaS* (SSU_RS02085) and *liaR* (SSU_RS02090), which have overlapping reading frames and are controlled by the same promoter. Thus we hypothesized that transposon insertions in *liaF* would disrupt the transcription of *liaS* and *liaR* and generated gene deletion mutants for *liaS* and *liaR* as a proxy for *liaF* gene deletion.

From the group of unknown genes of APS CEGs list, we selected SSU_RS09155, which had highest fold change (Table 1) and SSU_RS07155, which is predicted to encode a protein with unknown function and demonstrated a negative fold change in both APS and CSF. From the CEGs list in CSF, we also selected SSU_RS02635, encoding a protein with unknown function, and a fold change of -3,22 in CSF.

Site-specific gene deletion mutants were generated using our previously described CRISPR-Cas gene deletion method (Gussak et al.; see Methods) and verified by PCR amplification of the corresponding locus. Growth rates in THY and APS were assessed by OD_600_ measurements and CFU counting each hour during an 8-hour period (Figure 3). In THY medium, the mutant strains had the same growth rate as the wild type across all phases of the growth curve, except for the *liaR* mutant, which had a one-log^10^ reduced CFU count in the stationary phase (Figure 3). In APS, the Δ*purA* mutant exhibited growth attenuation compared to the wt, reaching a maximum concentration of approximately 10^7^ CFUs/ml, whereas the wt reached a concentration of nearly 10^9^ CFUs/ml. This growth defect correlated with the Tn-seq data, where *purA* had a log_2_FC difference of -3.19, indicating that functional PurA contributes to growth of *S. suis* P1/7 in APS. Growth assays in APS showed that Δ*liaR,* Δ*liaS* and ΔSSU_RS09155 gene deletion-mutants exhibited a growth delay during exponential phase and reached stationary phase with lower bacterial cell densities compared to the wild type, confirming the APS Tn-seq results (Figure 3). The *liaR* mutant did not exhibit any difference in growth in CSF compared to the wt parental strain, whereas in CSF, the Tn-seq results revealed a positive logFC for *liaR.* The ΔSSU_RS04755 mutant exhibited a delayed growth rate compared to the wt during the first 3 hours of growth in APS, although the mutant ultimately reached the same bacterial densities at stationary phase as wt. Mutants ΔSSU_RS07155 and ΔSSU_RS02635 did not show significantly altered growth curves in any of the tested conditions.

**Figure 3.**
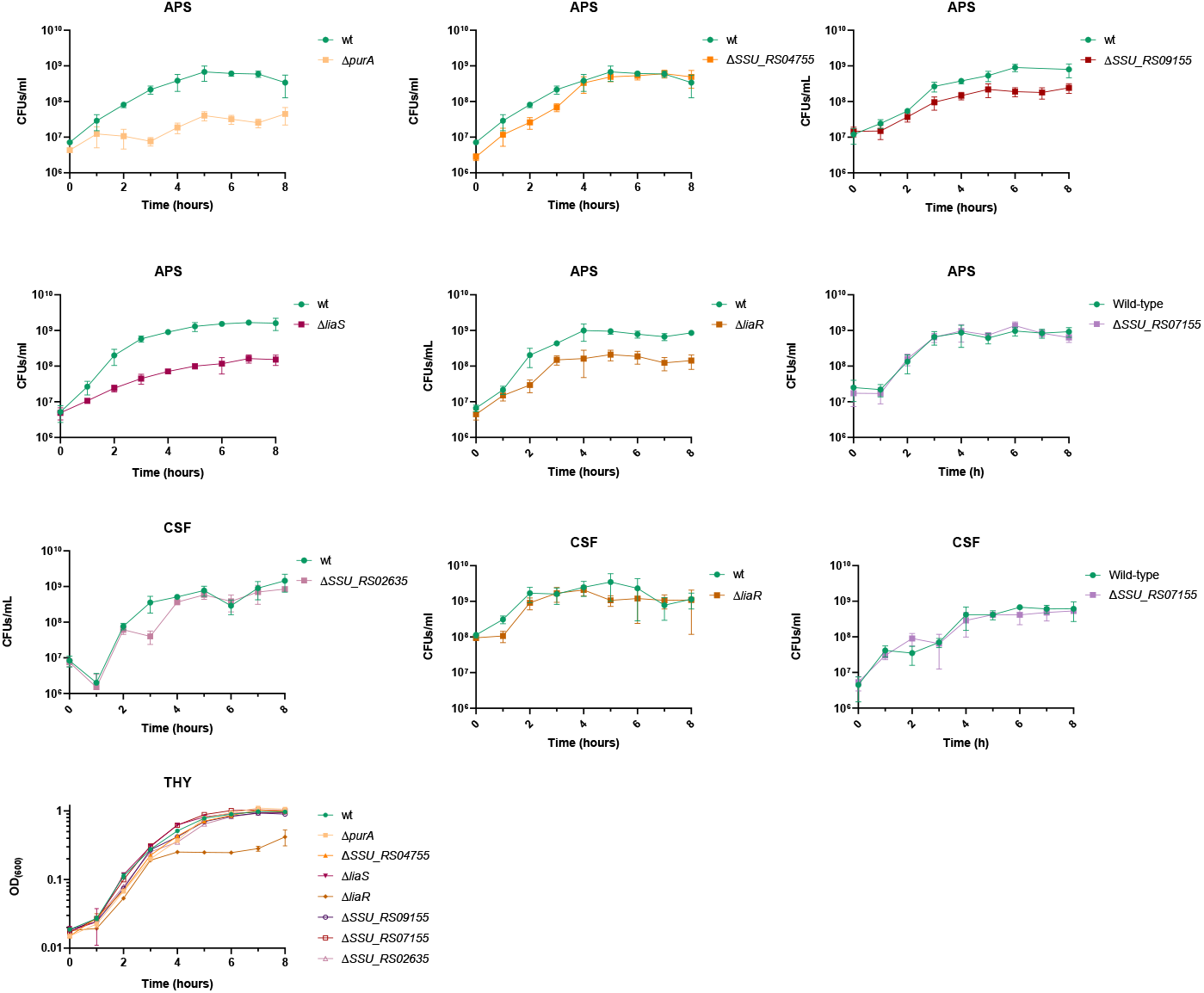
Growth curves in THY, APS and CSF. Data shown are means ± standard deviation of at least two biological replicates and two technical replicates for CFUs counting. CFUs/ml and OD_600_ axis are in Log_10_

### LiaR Activates Transcription of the Hypothetical Protein SSU_RS07195

Deletion of the gene encoding transcriptional regulator LiaR led to lower growth in APS. We hypothesized that LiaR may regulate a gene(s) critical for growth in APS. If our hypothesis were correct, we expected that some of the genes regulated by LiaR might feature in our Tn-seq results. To explore this, we used FIMO (Find Individual Motif Occurrences, part of the MEME online software suite, see Methods) to search for potential LiaR binding sites across the *S. suis* genome (Table S3). FIMO predicted presence of a putative LiaR binding site in the promoter of one of the genes encoding a hypothetical protein, SSU_RS07195, one of the genes identified as CEG in the APS Tn-seq results. To investigate possible functions of the hypothetical protein encoded by SSU_RS07195, we examined genomic position and neighbourhood of SSU_RS07195 in *S. suis* P1/7 in the NCBI database, and found the gene to be partially overlapping with gene SSU_RS07190, possibly part of a single operon together with a third gene SSU_RS10200 encoding an IS630 family transposase (Figure 4). The second gene SSU_RS07190 was predicted to encode a hypothetical protein containing a phage-shock protein (PspC) domain (Figure 4a). Both genes appear to be controlled by the same promoter, as the downstream gene lacks an independent promoter region, and the 5’ end of SSU_RS07190 overlaps with the end of SSU_RS07195. Both *liaF* and SSU_RS07195 are CEGs supporting growth of *S. suis* P1/7 in aps with a negative log_2_FC (−1.21 and -1.18, respectively; Table 1). To validate these predictions, we performed qPCR to determine whether expression of SSU_RS07195 was altered in the Δ*liaR* gene deletion mutant using *liaF* and *spx* genes, known to be regulated by LiaR in other species (Wolf et al., 2010) (Sanson et al., 2021) respectively, as references for comparison. Indeed, qPCR results showed that the expression of SSU_RS07195 was significantly reduced in *liaR* gene deletion mutant compared to wt, suggesting that LiaR positively regulates expression of SSU_RS07195 (Fig. 4). *liaF* and *spx* also showed reduced expression in *liaR* gene deletion mutant. Notably, the amount of downregulation of SSU_RS07195, -4.5 log_2_FC (Figure 4), was similar in magnitude to that observed for *liaF* and *spx*, -4.6 and -3.09 log_2_FC respectively (Figure 4).

**Figure 4.**
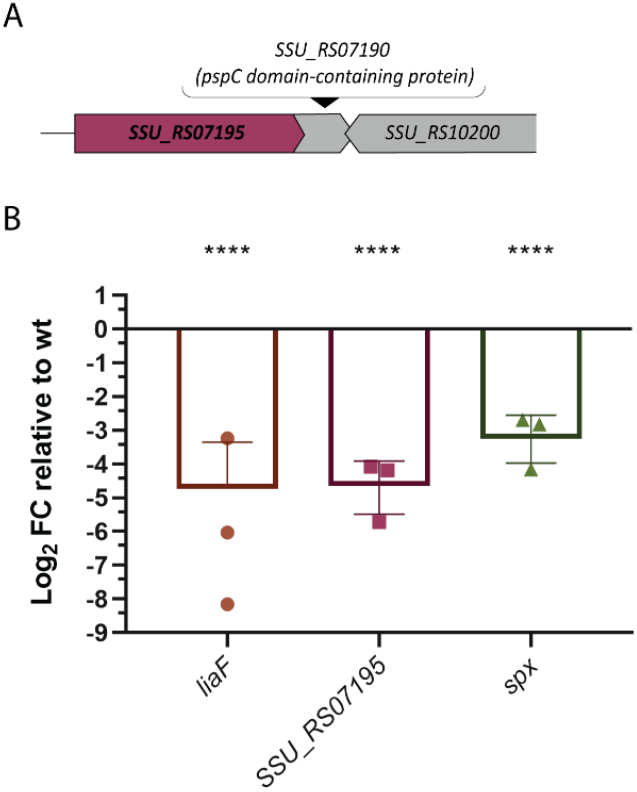
(A) Schematic representation of the SSU_RS07195 operon. (B) Expression of liaF, SSU_RS07195, and spx in the LiaR mutant relative to WT. Error bars represent standard deviation across biological replicates. Statistical analysis was performed using a one-way ANOVA (****, p < 0.0001).

### In silico analysis of *S. suis SSU_RS04755*

Gene SSU_RS04755 has been predicted to encode a basic membrane family protein (BMP), a transmembrane component of specific ABC transporters. Growth curve experiments did not reveal a significant reduction in the growth rate of our ΔSSU_RS04755 mutant. The operon genes with SSU_RS04755 were predicted to encode an ATP-binding protein and a permease which are typical components of an ABC transporter. Both predicted ATP-binding protein and permease encoding genes appeared as CEGs in our Tn-seq results (Table 1). The protein sequence associated with SSU_RS04755 shares significant identity with lipoproteins that are components of nucleoside ABC transporters in Gram-positive bacteria (see above), suggesting that SSU_RS04755 and its operon genes may encode an ABC transporter complex involved in nucleoside transport. To investigate possible functional equivalence of SSU_RS04755 operon genes with ABC transporters, we performed amino acid sequence alignments of SSU_RS04755 with PnrA and TmpC, which are homologous nucleoside binding lipoproteins from *S. pneumoniae* and *Treponema pallidum* respectively (Abdullah et al., 2021; Deka et al., 2006). Using the crystal structure of PnrA binding to adenosine (Abdullah et al., 2021), we confirmed that all but one (T70) of the specific amino acids involved in adenosine binding by PnrA were conserved in SSU_RS04755 (Figure 5c). To verify if the structural positions of these amino acids were conserved, we performed a structural prediction (see methods) of the predicted *S. suis* lipoprotein SSU_RS04755 and aligned it to the structure of PnrA (ID 6y9U in Protein Data Bank (PDB)). We confirmed that all amino acids involved in the adenosine binding are located at the same position in both protein structures. The non-conserved amino acid T70 which is I49 in *S. suis* might bind adenosine, as it has a carboxyl group (COOH) in the same position as the corresponding residue in PnrA that is involved in adenosine binding (Figure 5a and b).

**Figure 5.**
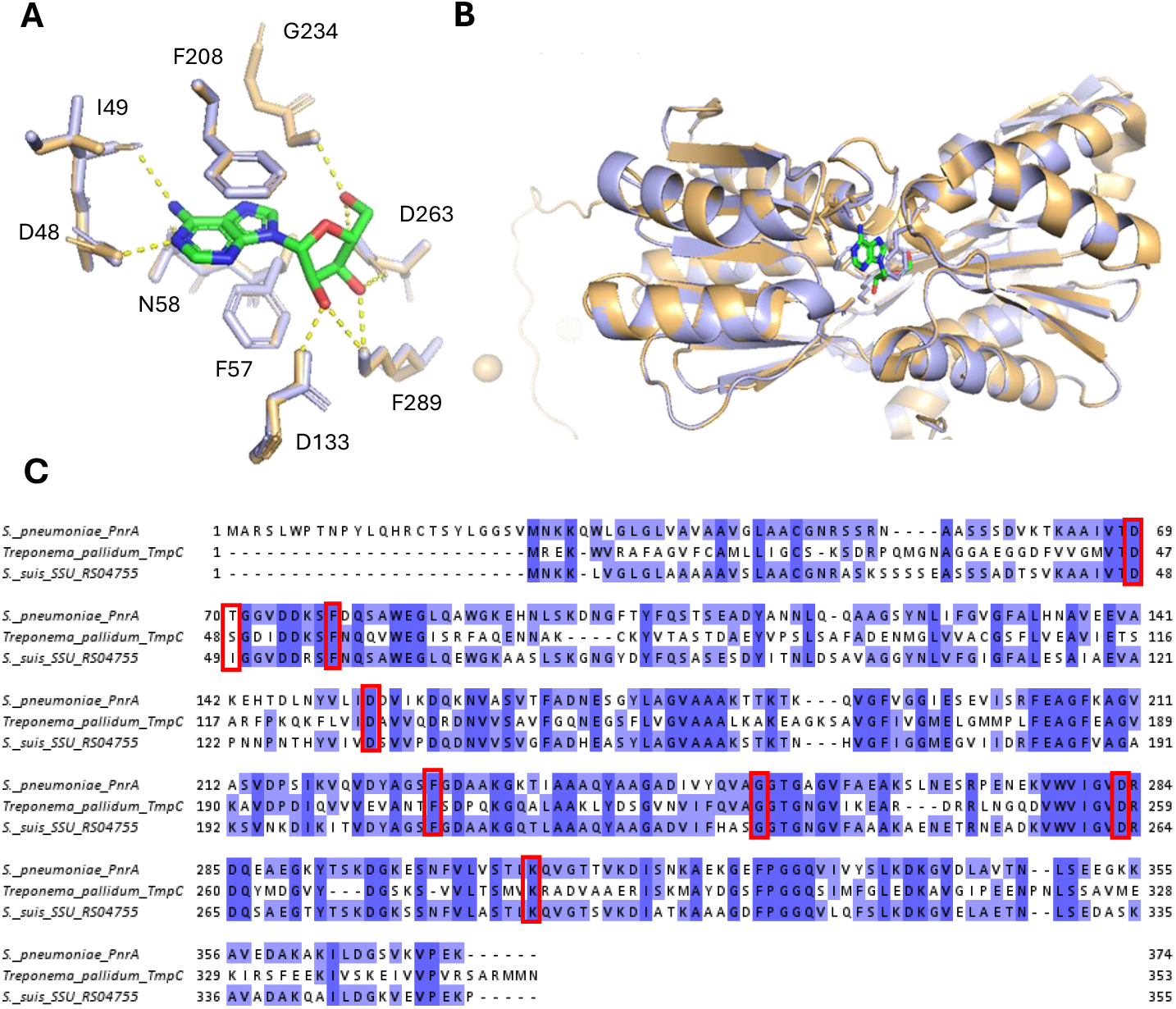
Structural and Sequence Analysis of Protein Homologs Featuring S. suis Lipoprotein SSU_RS04755 (blue) and S. pneumoniae PnrA (PDB ID: 6Y9U, Orange). (A) Zoomed-in view of the binding pocket with hydrogen bond interactions (dashed yellow lines). (B) Ribbon representation of the structural alignment of both proteins, highlighting active site interactions with a ligand (green) and key residues (sticks). (C) Multiple sequence alignment of homologous proteins from S. pneumoniae, Treponema pallidum, and S. suis, with conserved residues highlighted in blue and functionally significant residues marked with red boxes.

### Growth in CDM Reveals Nucleobase Uptake Deficiencies of Δ*SSU_RS04755* and Growth reduction of Δ*purA* Mutant

To further investigate our hypothesis that SSU_RS04755, SSU_RS04750, and SSU_RS04745 are part of an ABC transporter for nucleosides, we grew the ΔSSU_RS04755 mutant in CDM containing varying concentrations of purines and pyrimidines. When adenine was added to the CDM (Figure 6a), the ΔSSU_RS04755 mutant showed reduced growth compared to the wt, reaching an OD_600_ of approximately 0.4 versus 0.6 for the wt. Interestingly, the growth of both wt and ΔSSU_RS04755 with adenine supplementation was similar to their growth without nucleobases, suggesting a defect in the ΔSSU_RS04755 mutant’s ability to uptake adenine. We observed that wt grew the same with and without addition of nucleobases, which might suggest that during preculture in THY prior to CDM growth assays, wt bacteria stored nucleobases intracellularly. However, following the same preculturing step in THY, growth of the corresponding ΔSSU_RS04755 deletion mutant in CDM, with adenine as sole source of nucleobase, was substantially lower, in agreement with the hypothesis that SSU_RS04755 encodes a protein that is part of an ABC transporter importing nucleobases.

**Figure 6.**
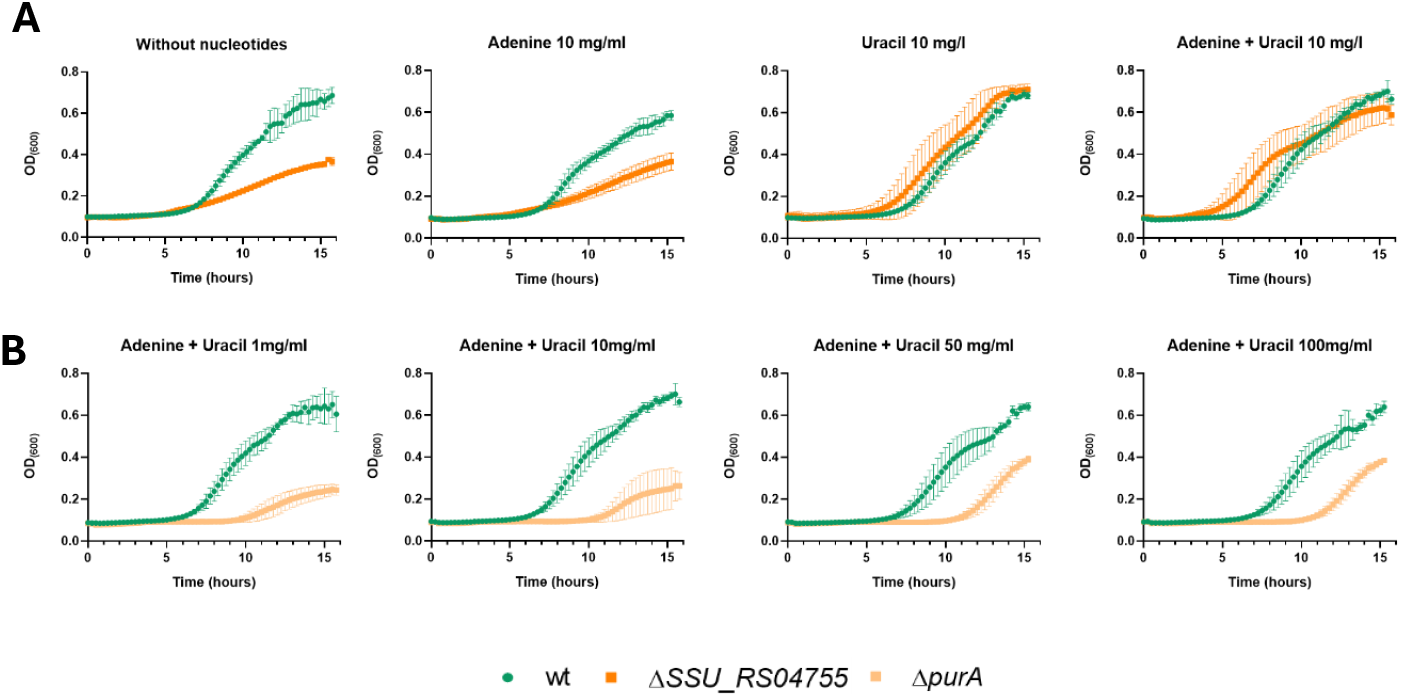
Growth curves of wt (green), purA (light orange) and ΔSSU_RS04755 (dark orange) in CDM. (A) wt and ΔSSU_RS04755 growth with different availability of adenine and uracil. (B) wt and ΔpurA growth with increasing concentrations of adenine and uracil. The experiments were performed with at least 3 biological replicates of each strain. Data shown are means ± standard deviation.

A second CEG, *purA,* was annotated to encode an adenylosuccinate synthase enzyme that is involved in the conversion of IMP (inosine monophosphate) to AMP (adenosine monophosphate). We hypothesized that the growth deficiency of the *purA* mutant in APS might have been due to its inability to synthesize adenine. We assessed growth of the *purA* mutant in CDM with increasing concentrations of nucleobases to compensate for the lack of de novo synthesized purines in the deletion mutant. Surprisingly, the addition of nucleobases could not recover the wt phenotype: the Δ*purA* deletion mutant exhibited a pronounced decrease in growth rate compared to the wt, reaching a maximum OD_600_ of approximately 0.4, whereas the wt OD_600_ reached up to 0.7. This phenotype was consistent regardless of the concentration of nucleobases added to the CDM (Figure 6b).

## Discussion

In this study, we constructed a transposon mutant library (Tn-library) for *S. suis* strain P1/7 and employed transposon sequencing (Tn-seq) to screen for conditionally essential genes (CEGs) that mediated growth of *S. suis* bacteria in two host body fluids, activated porcine serum (APS) and cerebrospinal fluid (CSF). APS was selected to simulate bloodstream conditions, as it supports more reproducible *S. suis* growth than whole blood, which likely restricts growth through neutrophil activation and antimicrobial peptide release (Bleuzé et al., 2024). Additionally, we chose to perform Tn-seq in CSF since virulent *S. suis* strains like P1/7 are able to grow in CSF. Using APS and CSF allowed controlled library screening while minimizing stochastic selection of mutants typical of *in vivo* models (Bourgeois & Camilli, 2023). To our knowledge, this is the first study to apply an in-house Nanopore sequencing approach to Tn-seq in *S. suis.* For downstream analysis, we designed a Tn-Seq data analysis pipeline to analyze nanopore sequencing output and generate a list of CEGs.

The Tn-seq results revealed a list of 33 and 25 CEGs that appeared crucial for optimal *S. suis* growth in APS and CSF, respectively. According to the functional annotations of the crucial genes mediating growth in APS, the main biological processes regulated by these genes were *de novo* biosynthesis of nucleotides and ABC transport systems. Genes conditionally essential for growth in CSF were primarily involved in amino acid transport. Five genes were common to both lists; three of these were part of a tryptophan ABC transporter operon, which has been previously reported as conditionally essential in Tn-seq studies on *S. suis*, corroborating reports that *S. suis* can not synthesize tryptophan (Arenas et al., 2020; Dresen et al., 2022). In APS, a substantial number of genes were annotated to be involved in nucleotide metabolism and nucleotide transport. Arenas and colleagues and Dresen and colleagues studied genes supporting *S. suis* growth in blood and CSF using a porcine *in vivo* Tn-library approach, and found that genes involved in *de novo* nucleotide biosynthesis, such as *purA, purB*, and *guaA*, were conditionally essential in both fluids (Arenas et al., 2020; Dresen et al., 2022). The genes *purA* and *guaB* were also reported as CEGs for growth of Group A Streptococcus (GAS) in human blood (Breton et al., 2013), suggesting that genes related to nucleotide metabolism and transport are important for survival of streptococci in APS. Our Tn-seq screening in CSF did not identify CEGs annotated with functions in nucleotide biosynthesis. However, it revealed an enrichment of 10 genes involved in protein synthesis, 8 of which were annotated with functions in amino acid transport systems. This finding is consistent with reports showing that CSF contains lower amino acid levels than serum (Otto et al., 2024), which may necessitate the upregulation of amino acid transport systems for growth. Notably, these same studies identified inosine, a purine nucleoside that plays an essential role in purine nucleotide biosynthesis, as one of the metabolites found exclusively in CSF (Otto et al., 2024). This may explain the absence of CEGs annotated with functions in nucleotide transport and biosynthesis pathways in the Tn-seq results from CSF, as inosine in CSF could act as a readily available precursor for purine nucleotide synthesis.

From our list of CEG in APS, five genes, i.e. *purA*, SSU_RS04755, SSU_RS04755, *liaF*, SSU_RS09155 and SSU_RS07155 were selected for verification of the library results by generating gene-specific in-frame deletion mutants. From our list of CEG in CSF, genes SSU_RS02635 and SSU_RS07155 with negative FC and *liaR* with positive FC were selected for verification. The choice of these genes was based on: (i) enrichment of certain biological processes according to CEG annotations; (ii) genes identified in Tn-library based studies in different streptococci, i.e. *purA* and *liaF*; (iii) uncharacterized genes in *S. suis* of possible interest, f.e. located in operon with gene of interest; and (iv) presence of a CEG in APS and CSF. We failed in obtaining viable *liaF* deletion mutants, but could obtain deletion mutants of operon genes *liaS* and *liaR*. Deletion mutants of the genes *purA, liaR, liaS* and SSU_RS09155 had lower growth rates in APS and reached lower CFU/ml counts in stationary phase compared to the wt strain, thus validating the Tn-seq results. The *liaR* deletion mutant showed no growth deficiency in CSF compared to the wt, thus confirming the results obtained for this gene in CSF. Gene deletion mutants for the genes SSU_RS04755, SSU_RS02635, and SSU_RS07155 did not exhibit any significant reduction in growth rate under any of the conditions tested. This could be due to competition among mutants during library growth or a polar effect of the transposon insertion on downstream genes.

*In silico* analysis suggested that gene SSU_RS04755 encodes a lipoprotein that is part of a nucleotide ABC transporter. Notably, all three genes in the SSU_RS04745–55 operon were identified as CEGs in APS, underscoring the importance of this nucleotide ABC transporter for survival in serum-like conditions, highlighting the importance of this candidate nucleotide ABC transporter to support *S. suis* growth in APS. Our growth experiments in CDM demonstrated that the SSU_RS04755 gene deletion mutant showed significant growth reduction when purines were the sole source of nucleobases, suggesting its essential role in purine uptake. Taken together, these results support the *in silico* predictions that the SSU_RS04745-55 operon encodes an ABC transporter for purine uptake. In *S. pneumoniae* a nucleoside transporter with 66% homology to the *S. suis* SSU_RS04745-55 candidate purine transporter was shown to be important for growth in serum. Additionally, *S. pneumoniae* Δspd_0739 mutant, which encodes a lipoprotein that is part of a nucleotide transport system, exhibited reduced virulence compared to wild-type (Saxena et al., 2015). The spd_0739 lipoprotein component of the transporter was proposed as a vaccine candidate due to its high conservation among strains (>98%), its expression both *in vitro* and during *in vivo* infection, and its low homology (<11%) with human lipoproteins (Saxena et al., 2015). Additionally, in *Borrelia burgdorferi*, two purine uptake transporters have been reported as important for *in vivo* infection in mice, and mutants lacking these purine transporters showed no impaired growth in rich media *in vitro* (Jain et al., 2012) as we observed in our *in vitro* results. These results highlight the importance of purine transport and acquisition for in-host survival and pathogenesis.

We identified the genes *purA, purB, guaA, guaB* and *pyrE* involved in *de novo* nucleotide biosynthesis as CEGs supporting *S. suis* growth in APS. Previous studies had shown that disruption in the purine biosynthesis pathways may reduce bacterial colonization and intracellular growth, increase susceptibility to oxidative stress, and lower bacterial proliferation in human serum and blood (Li et al., 2018; Mantena et al., 2008; Samant et al., n.d.; Schauer et al., 2010). Additionally, genome-wide screens in *Staphylococcus aureus* demonstrated that *pur* and *pyr* gene deletion mutants were attenuated in various infection models (Lan et al., 2010; Mei et al., 1997; Valentino et al., 2014; Wilde et al., 2015). The importance of the nucleotide biosynthesis pathway is not surprising, as it is (i) essential for the synthesis of nucleic acids; (ii) mediating production of signaling molecules like cyclic AMP and GMP and (iii) generating energy storage molecules such as ATP and GTP (Goncheva et al., 2022). Our *S. suis* P1/7 *purA* mutant showed significant growth defects in CDM and was not rescued by the addition of nucleobases suggesting that in limited-nutrient media, nucleobases uptake could not rescue viability-lowering effects of *purA* deletion. It thus appears that nucleobase uptake and salvage pathways alone are insufficient to meet the purine demands of the *S. suis* P1/7Δ*purA* mutant. However, while several studies in other bacterial genera such as *Escherichia, Salmonella*, and *Bacillus* have demonstrated successful growth restoration of *pur* and *pyr* mutants following the addition of nucleobases to the media (Samant et al., n.d.; Shaffer et al., 2017), no studies have been found regarding growth restoration in *Streptococcus*. Given that *de novo* purine biosynthesis involves an extensive list of *pur* genes, it appears that mutations at different points in the pathway may produce distinct phenotypic effects such that compensatory mutations or pathway rerouting is usually not restoring viability nor growth recovery. In studies where growth recovery had turned out possible, deleted genes were typically at the beginning of the pathway, such as *purF* and *purE* (Samant et al., n.d.; Shaffer et al., 2017). In contrast, in *S. suis* P1/7, *purA* is located toward the end of the pathway (KEGG. *Purine metabolism – Pathway ID: ssi00230)* and may therefore result in more pronounced phenotypic changes. The significance of *purA* in pathobiology is further highlighted by a study in clinical *S. aureus*, where it was found that *purA* was induced when *S. aureus* entered into a semi-dormant state in the human body upon exposure to acidic pH or neutrophils as part of the immune response to *S. aureus* infection (Huemer et al., 2020).

Our study revealed the important regulatory role of the LiaFSR three-component system in mediating *S. suis* growth in APS, with a particular focus on the regulatory function of LiaR. Our data suggest that LiaR acts as a positive regulator of at least one CEG involved in supporting *S. suis* growth in APS, highlighting its role in stress adaptation and environmental sensing. Of note, *liaF* and gene SSU_RS07195 encoding a hypothetical protein displayed negative FC values during growth in APS, but positive FC values during growth in CSF, implying that LiaFSR activation is advantageous in APS, while its repression may be beneficial or neutral in CSF. This key role of LiaFSR sensory system to adapt to different host niches and nutrient availability has been described previously by Sanson and colleagues (Sanson et al., 2021) who reported that in *Streptococcus pyogenes*, a mutant not producing LiaR showed reduced expression of virulence genes including *spxA2*, which is involved in oxidative stress response, in an *ex vivo* human blood model. We found that our *S. suis* P1/7 *liaR* gene deletion mutant exhibited reduced growth during culture in APS, a model for growth in blood (*S. suis* P1/7 cannot be cultured reproducibly in whole blood) while the deletion of S. suis P1/7 *liaR* did not reduce bacterial growth during culture in CSF; instead, our Tn-seq data suggested that deletion of *liaR* was associated with increased bacterial growth in CSF. These results highlight the metabolic flexibility that disease-associated bacteria can be capable of in order to rapidly alter gene expression to adapt to different host niches. These results also reinforce the notion that some candidate antibacterial targets are only expressed in specific host niches, so that treatments including vaccinations should take such niche-specific antibacterial target expression into account when treating bacterial infections.

## Data Availability

The raw nanopore sequencing data generated in this study are currently available from the corresponding author upon reasonable request. The data will be deposited in a public repository, and accession details will be provided upon publication.

### Acknowledgments

We thank Tim Van Opijnen’s lab (Tufts University) for the transposon library protocol and plasmids, and Alex Gussak (Wageningen University & Research) for the CRISPR-Cas9 protocol for *S. suis* mutants design.

