## Supplemental Table 1 for "Genome-Wide Identification of Conditionally Essential Genes Supporting *Streptococcus suis* Growth in Serum and Cerebrospinal Fluid"

Table S1.Stock materials for CDM

| | |
|---|---|
| **Composition of CDM buffer in 100 ml water** | |
| Na-B-glycerophosphate | 9.3 g |
| $KH_2PO_4$ | 0.4 g |
| $(NH_4)_2$ citrate | 0.25 g |
| Na acetate | 0.44 g |
| **Adjust pH to 6.4* | |
| Bacto tryptone | 7.5 g |
| **Metal mixture in 10 ml water** | |
| $MgCl_2$ | 0.103 g |
| $CaCl_2.2H_2O$ | 54 mg |
| $ZnSO_4.7H_2O$ | 5.5 mg |
| $CoSO_4.7H_2O$ | 3.3 mg |
| $CuSO_4.5H_2O$ | 0.17 mg |
| **Vitamin mixture in 10 ml water** | |
| Pyridoxal-Cl | 2 mg |
| Thiamine $Cl_2$ | 1 mg |
| Riboflavin | 1 mg |
| Ca-pantothenate | 1 mg |
| Biotin | 0.1 mg |
| Folic acid | 1 mg |
| Vitamin B6 | 1 mg |
| **Light sensitive reagents, adjust pH to 7* | |
| **Amino acids** | **Concentration (g/L)** |
| Alanine | 3 |
| Glycine | 3 |
| Arginine | 3 |
| Serine | 3 |
| Threonine | 6 |
| Cysteine | 3 |
| Proline | 3 |
| Asparagine | 3 |
| Aspartate | 3 |
| Methionine | 3 |
| Lysine | 3 |
| Glutamine | 6 |
| Histidine | 3 |
| Glutamate | 3 |
| Phenylalanine | 3 |
| Tryptophan | 3 |
| Valine | 3 |
| Leucine | 3 |
| Isoleucine | 3 |
| **Additional stock material** | **Concentration** |
| $MnSO_4.H_2O$ | 28 mg/ml |
| Choline chloride | 2.5 mg/ml |
| Pyruvate | 1 mg/ml |
| Glucose | 50 M |

| **Nucleotide/ nucleoside** | **Concentration (mg/ml)** |
|---|---|
| Adenine | 1 |
| Uracil | 1 |
| Thymidine | 1 |
| Cytidine | 1 |
| Guanine | 1 |
