## Supplemental Table 2 for "Genome-Wide Identification of Conditionally Essential Genes Supporting *Streptococcus suis* Growth in Serum and Cerebrospinal Fluid"

Table S2. Primers list

| Primer | Sequence | Used for |
|---|---|---|
| P001 | tagtaccaaccttcaaatgattcccacggctcttttatacgtaaaggac | Chl_smal_Fwd |
| P002 | ggatccaatttttttataattttttttaatctgttcctcgagtctagaactaagaatggc | Chl_swal_Rev |
| P003 | ttccctacacgacgctcttccgatctatcacgnn | Adapter_Fwd |
| P004 | /5phos/cgtgatagatcggaagagcgtcgtgtagggaaagagt | Adapter_Rev |
| P005 | caagcagaagacggcatacgaagaccggggacttatcatccaacctgt | HimarTn_Fwd |
| P006 | aatgatacggcgaccaccgagatctacactctttccctacacgacgctcttccgatct | Adpt_enrich_Rev |
| P007 | tgatcaaacaagacacgcttgcct | Guide1_fwd_ purA |
| P008 | aaacaggcaagcgtgtcttgtttg | Guide1_rev_ purA |
| P009 | tgatgctccaaacttgctgggtag | Guide2_fwd_ purA |
| P010 | aaacctacccagcaagtttggagc | Guide2_rev_ purA |
| P011 | tgatgcagccttgtccatgtaggc | Guide3_fwd_ purA |
| P012 | aaacgcctacatggacaaggctgc | Guide3_rev_ purA |
| P013 | tgatattgaagtacttgaccgcca | Guide1_fwd_SSU_RS04755 |
| P014 | aaactggcggtcaagtacttcaat | Guide1_rev_SSU_RS04755 |
| P015 | tgattaccagtaccaccagaagcg | Guide2_fwd_SSU_RS04755 |
| P016 | aaaccgcttctggtggtactggta | Guide2_rev_SSU_RS04755 |
| P017 | tgatgaagaccttcccaagcaga | Guide3_fwd_SSU_RS04755 |
| P018 | aaacatctgcttgggaaggtcttc | Guide3_rev_SSU_RS04755 |
| P019 | tgatagaacggaaacggattgcca | Guide1_fwd_liaS |
| P020 | aaactggcaatccgtttccgttct | Guide1_rev_liaS |
| P021 | tgatcagttgaaaatgacagacga | Guide2_fwd_liaS |
| P022 | aaactcgtctgtcattttcaactg | Guide2_rev_liaS |
| P023 | tgattgtcagcctagcctccactc | Guide3_fwd_liaS |
| P024 | aaacgagtggaggctaggctgaca | Guide3_rev_liaS |
| P025 | tgatgtgctggaagcaggagctcg | Guide1_fwd_liaR |
| P026 | aaaccgagctcctgcttccagcac | Guide1_rev_liaR |
| P027 | tgataatcttagcagccatccgca | Guide2_fwd_liaR |
| P028 | aaactgcggatggctgctaagatt | Guide2_rev_liaR |
| P029 | tgatatcttgacgctgttagccaa | Guide3_fwd_liaR |
| P030 | aaacttggctaacagcgtcaagat | Guide3_rev_liaR |
| P031 | tgatcatcgatatgaaaatctgcc | Guide1_fwd_SSU_RS09155 |
| P032 | aaacggcagattttcatatcgatg | Guide1_rev_SSU_RS09155 |
| P033 | tgatcatggatgtatcgagtcgag | Guide2_fwd_SSU_RS09155 |
| P034 | aaacctcgactcgatacatccatg | Guide2_rev_SSU_RS09155 |
| P035 | tgatggttaatacaaaccagacag | Guide3_fwd_SSU_RS09155 |
| P036 | aaacctgtctggtttgtattaacc | Guide3_rev_SSU_RS09155 |
| P037 | tgattcttctgtcaaatccactgt | Guide1_fwd_SSU_RS07155 |
| P038 | aaacacagtggatttgacagaaga | Guide1_rev_SSU_RS07155 |
| P039 | tgatgatttgctcaaaggtttcca | Guide2_fwd_SSU_RS07155 |
| P040 | aaactggaaacctttgagcaaatc | Guide2_rev_SSU_RS07155 |
| P041 | tgatagcagctgcagcagctccgc | Guide3_fwd_SSU_RS07155 |
| P042 | aaacgcggagctgctgcagctgct | Guide3_rev_SSU_RS07155 |
| P043 | tgatgtgaccattcctacaccgtt | Guide1_fwd_SSU_RS02635 |
| P044 | aaacaacggtgtaggaatggtcac | Guide1_rev_SSU_RS02635 |
| P045 | tgatagtggatcagcaatttcaag | Guide2_fwd_SSU_RS02635 |
| P046 | aaaccttgaaattgctgatccact | Guide2_rev_SSU_RS02635 |
| P047 | tgattgatctggacgggcagattg | Guide3_fwd_SSU_RS02635 |

| | | |
|---|---|---|
| P048 | aaaccaatctgcccgtccagatca | Guide3_rev_SSU_RS02635 |
| P049 | ggcatggagttcaggaacatac | HA1_fwd_purA |
| P050 | tggatgaattgccagaagcg | HA1_rev_purA |
| P051 | cgcttctggcaattcatccacaagtgcaatttatacttcgtgcc | HA2_fwd_purA |
| P052 | gaggaagttttgtgctaatttgagg | HA2_rev_purA |
| P053 | gggtcaatacagctgtcgg | HA1_fwd_ SSU_RS04755 |
| P054 | gcagctgcaggtgatttcc | HA1_rev_ SSU_RS04755 |
| P055 | ggaaatcacctgcagctgcacatgcagccaaagatacagc | HA2_fwd_ SSU_RS04755 |
| P056 | gttgctttagcagctgaaagc | HA2_rev_ SSU_RS04755 |
| P057 | ctatcttcgacttcgtttcctagg | HA1_fwd_ liaS |
| P058 | gtgggatacgaatatcgatcgcaagtgggagggtggaagc | HA1_rev_ liaS |
| P059 | gcgatcgatattcgtatcccac | HA2_fwd_ liaS |
| P060 | ctgatggtacactacctgacc | HA2_rev_ liaS |
| P061 | cattgctagatgttacgctctgg | HA1_fwd_ liaR |
| P062 | ttctgcaactacttcaacatctgg | HA1_rev_ liaR |
| P063 | ccagatgttgaagtagttgcagaagctgttagcgatcgtactcaag | HA2_fwd_ liaR |
| P064 | ttgatatgcgcaactggtcc | HA2_rev_ liaR |
| P065 | agcaacattccgacaaaaacg | HA1_fwd_ SSU_RS09155 |
| P066 | acgggtgatatggcattttctac | HA1_rev_ SSU_RS09155 |
| P067 | gtagaaaatgccatatcacccgtcagtcctataactgcagctcc | HA2_fwd_ SSU_RS09155 |
| P068 | ctgcccagtataaaatggaac | HA2_rev_ SSU_RS09155 |
| P069 | cggtctgctcaattagacgg | HA1_fwd_ SSU_RS07155 |
| P070 | atcgaagatgttcaaactgaagtgg | HA1_rev_ SSU_RS07155 |
| P071 | ccacttcagtttgaacatcttcgatcaaaaggctagtccacacgc | HA2_fwd_ SSU_RS07155 |
| P072 | acggtggtattgccatctatgg | HA2_rev_ SSU_RS07155 |
| P073 | agccctgattctttggatgc | HA1_fwd_ SSU_RS02635 |
| P074 | ccagatcaatttcatttcg | HA1_rev_ SSU_RS02635 |
| P075 | cgaaatgaaattgatctggccagcagctttttgcttcc | HA2_fwd_ SSU_RS02635 |
| P076 | acgctttctcacagagaaagg | HA2_rev_ SSU_RS02635 |
| P077 | cgcattgatttgagtcagctagg | Check_guide_pSStarget_fwd |
| P078 | tcggtgccactttttcaagttg | Check_guide_pSStarget_rev |
| P079 | gcgtgggtcagcatatttacg | qPCR_fwd_gyrA |
| P080 | cgtcttcgtcgtttgacagg | qPCR_rev_gyrA |
| P081 | ccacgaccactctcaagatagg | qPCR_fwd_liaF |
| P082 | gatactagttgctgaaagtgcaacc | qPCR_rev_liaF |
| P083 | ccaagacatgacctgattgcg | qPCR_fwd_SSU_RS07195 |
| P084 | ggatattcgcttgacgacttgc | qPCR_rev_ SSU_RS07195 |
| P085 | tgaagagttgagtgtgaacgag | qPCR_fwd_spx |
| P086 | tgagcattttcaacattgcgc | qPCR_rev_ spx |
