## Supplemental Table 3 for "Genome-Wide Identification of Conditionally Essential Genes Supporting *Streptococcus suis* Growth in Serum and Cerebrospinal Fluid"

Table S3. FIMO results

| motif_alt_id | start | stop | strand | p-value | matched_sequence | Gene_id | |
|---|---|---|---|---|---|---|---|
| **THMDTCNWMAGWYYKA** | 1455682 | 1455697 | - | 2.89E-07 | TCAGTCCTAAGACTGA | SSU_RS07275 | |
| **THMDTCNWMAGWYYKA** | 586897 | 586912 | - | 6.13E-07 | TAATTCCTCAGACTGA | SSU_RS02955 | |
| **THMDTCNWMAGWYYKA** | 133852 | 133867 | - | 2.83E-06 | TCCATCTTGAGACCGA | SSU_RS00875 | |
| **THMDTCNWMAGWYYKA** | 1471430 | 1471445 | + | 2.83E-06 | TAGGTCTACAGACCGA | SSU_RS07360 | |
| **THMDTCNWMAGWYYKA** | 966661 | 966676 | + | 4.25E-06 | TTCTTCATAAGGCCGA | SSU_RS04785 | |
| **THMDTCNWMAGWYYKA** | 1200282 | 1200297 | + | 4.89E-06 | TAGGTCTACAGACCTA | SSU_RS05975 | |
| **THMDTCNWMAGWYYKA** | 78685 | 78700 | - | 1.25E-05 | TTCTTCGTCAGTGTTA | SSU_RS00495 | |
| **THMDTCNWMAGWYYKA** | 1371777 | 1371792 | + | 1.25E-05 | TCCTTCCACGGTCTTA | SSU_RS06795 | |
| **THMDTCNWMAGWYYKA** | 66354 | 66369 | - | 1.72E-05 | TCATTCAACAGACCTG | SSU_RS00385 | |
| **THMDTCNWMAGWYYKA** | 761780 | 761795 | - | 1.72E-05 | ATAGTCAACAGTCCTA | SSU_RS03750 | |
| **THMDTCNWMAGWYYKA** | 914888 | 914903 | + | 1.72E-05 | TTTATCATAAGTTCGA | SSU_RS04510 | |
| **THMDTCNWMAGWYYKA** | 1200266 | 1200281 | - | 1.72E-05 | TTCTTCAATAGTTTGA | SSU_RS05975 | |
| **THMDTCNWMAGWYYKA** | 78562 | 78577 | - | 1.86E-05 | TCCGTCAAATGACTTA | SSU_RS00495 | |
| **THMDTCNWMAGWYYKA** | 1455682 | 1455697 | + | 1.86E-05 | TCAGTCTTAGGACTGA | SSU_RS07275 | |
| **THMDTCNWMAGWYYKA** | 1631626 | 1631641 | - | 2.23E-05 | ACCATCGAAAGTTTGA | SSU_RS08165 | |
| **THMDTCNWMAGWYYKA** | 1222959 | 1222974 | - | 2.55E-05 | TACATCTAAAGTCCTC | SSU_RS06075 | |
| **THMDTCNWMAGWYYKA** | 85202 | 85217 | - | 2.85E-05 | TCCATCAAACGTTTGA | SSU_RS00565 | |
| **THMDTCNWMAGWYYKA** | 1380224 | 1380239 | + | 3.20E-05 | TACTTCGAAAGTTTGT | SSU_RS06845 | |
| **THMDTCNWMAGWYYKA** | 1432912 | 1432927 | - | 3.20E-05 | TTATTCGAAAGTCTGG | SSU_RS07135 | |
| **THMDTCNWMAGWYYKA** | 75003 | 75018 | + | 3.52E-05 | TACGTCGAAACTTCTA | SSU_RS00450 | |
| **THMDTCNWMAGWYYKA** | 914860 | 914875 | - | 3.52E-05 | TCAATCGAAATACTGA | SSU_RS04510 | |
| **THMDTCNWMAGWYYKA** | 617409 | 617424 | - | 3.76E-05 | TAAATCAAAGGTCCTA | SSU_RS03085 | |
| **THMDTCNWMAGWYYKA** | 950549 | 950564 | - | 3.76E-05 | AAAATCAACAGATTGA | SSU_RS04700 | |
| **THMDTCNWMAGWYYKA** | 1455675 | 1455690 | + | 3.76E-05 | CTATTCATCAGTCTTA | SSU_RS07275 | |
| **THMDTCNWMAGWYYKA** | 1123024 | 1123039 | - | 4.30E-05 | TAAATCGACGGTTTTA | SSU_RS05595 | |
| **THMDTCNWMAGWYYKA** | 744487 | 744502 | + | 4.53E-05 | TTAGTCCAAAATCTTA | SSU_RS03655 | |
| **THMDTCNWMAGWYYKA** | 471962 | 471977 | + | 4.53E-05 | TCAGTCGAAAAATTTA | SSU_RS02380 | |
| **THMDTCNWMAGWYYKA** | 180772 | 180787 | + | 4.70E-05 | TTCATCATAAGATTTT | SSU_RS01070 | |
| **THMDTCNWMAGWYYKA** | 420081 | 420096 | - | 4.91E-05 | TTCATGAAAAGTTCTA | SSU_RS02125 | |
| **THMDTCNWMAGWYYKA** | 1391045 | 1391060 | - | 5.13E-05 | TACATCGTAAAATTTA | SSU_RS06895 | |
| **THMDTCNWMAGWYYKA** | 277657 | 277672 | - | 5.33E-05 | TTATTCTTCATTTTTA | SSU_RS01470 | |
| **THMDTCNWMAGWYYKA** | 427319 | 427334 | + | 5.33E-05 | TAATTTATAAGTTTGA | SSU_RS10460 | |
| **THMDTCNWMAGWYYKA** | **952730** | **952745** | **+** | **5.57E-05** | **TATCTCCTCAGTCCTA** | **SSU_RS04710** | **spx** |
| **THMDTCNWMAGWYYKA** | **68995** | **69010** | **+** | **5.61E-05** | **TAGGTCTGCAGACCGA** | **SSU_RS00395** | **spx** |
| **THMDTCNWMAGWYYKA** | 68995 | 69010 | - | 5.63E-05 | TCGGTCTGCAGACCTA | SSU_RS00395 | |
| **THMDTCNWMAGWYYKA** | 1465466 | 1465481 | + | 6.25E-05 | TCAGTCTCTAGACCTA | SSU_RS07330 | |
| **THMDTCNWMAGWYYKA** | **412250** | **412265** | **-** | **6.70E-05** | **TCCATCCAAAGTAGGA** | **SSU_RS02080** | **liaF** |
| **THMDTCNWMAGWYYKA** | 1929398 | 1929413 | + | 6.70E-05 | TCCGTCGCATGATCGA | SSU_RS09535 | |
| **THMDTCNWMAGWYYKA** | 165544 | 165559 | - | 6.95E-05 | TTCCTCAACAGCTTTA | SSU_RS01005 | |
| **THMDTCNWMAGWYYKA** | 1959339 | 1959354 | + | 6.95E-05 | TTCCTCAATAGTTTGA | SSU_RS09680 | |
| **THMDTCNWMAGWYYKA** | 813194 | 813209 | - | 7.08E-05 | GGCTTCGAAAGTCTGA | SSU_RS04000 | |
| **THMDTCNWMAGWYYKA** | 1465466 | 1465481 | - | 7.89E-05 | TAGGTCTAGAGACTGA | SSU_RS07330 | |
| **THMDTCNWMAGWYYKA** | 704128 | 704143 | - | 8.09E-05 | TTCTTCCTCTGACCAA | SSU_RS03495 | |
| **THMDTCNWMAGWYYKA** | 1393059 | 1393074 | + | 8.78E-05 | TTCTTCCACAGCTATA | SSU_RS06910 | |
| **THMDTCNWMAGWYYKA** | 1785548 | 1785563 | + | 8.78E-05 | TCCTTCCAAAGACAGG | SSU_RS08865 | |
| **THMDTCNWMAGWYYKA** | 1095781 | 1095796 | - | 9.15E-05 | GCAGTCGTCAGCTTGA | SSU_RS05480 | |
| **THMDTCNWMAGWYYKA** | **1442378** | **1442393** | **+** | **9.15E-05** | **TCAGCCCGAAGACTGA** | **SSU_RS07195** | |
| **THMDTCNWMAGWYYKA** | 389373 | 389388 | - | 9.61E-05 | TGCTTCATAAAACCGA | SSU_RS01980 | |
